# Integration of transcriptomics and network analysis reveals co-expressed genes in *Frankliniella occidentalis* guts that respond to tomato spotted wilt virus infection

**DOI:** 10.1101/2021.08.14.456355

**Authors:** Jinlong Han, Dorith Rotenberg

## Abstract

**Background:** The arthropod gut is the first barrier to infection by viruses that are internally borne and transmitted persistently by arthropod vectors to plant and animal hosts. Tomato spotted wilt virus (TSWV), a plant-pathogenic virus, is transmitted exclusively by thrips vectors in a circulative-propagative manner. *Frankliniella occidentalis* (western flower thrips), the principal thrips vector of TSWV, is transmission-competent only if the virus is acquired by young larvae. To begin to understand the larval gut response to TSWV infection and accumulation, a genome-assisted, transcriptomic analysis of *F. occidentalis* gut tissues of first (early L1) and second (early L2 and late L2) instar larvae was conducted using RNA-Seq to identify differentially-expressed transcripts (DETs) in response to TSWV compared to non- exposed cohorts.

**Results:** The larval gut responded in a developmental stage-dependent manner, with the majority of DETs (71%) associated with the early L1 stage at a time when virus infection is limited to the midgut epithelium. Provisional annotations of these DETs inferred roles in digestion and absorption, insect innate immunity, and detoxification. Weighted gene co-expression network analysis using all assembled transcripts of the gut transcriptome revealed eight gene modules that distinguish the larval development. Intra-module interaction network analysis of three most DET- enriched modules revealed ten central hub genes. Droplet digital PCR-expression analyses of select network hub and connecting genes revealed temporally-dynamic changes in gut expression during and post exposure to TSWV.

**Conclusion:** These findings expand our understanding of the developmentally-mediated interaction between thrips vectors and orthotospoviruses, and provide opportunities for probing pathways for biomarkers of thrips vector competence.

## BACKGROUND

The alimentary canal of arthropods is the primary site of entry into vectors by various viral pathogens because of its constant interactions with surrounding environment by insect feeding [1, 2]. In contrast to externally-borne non-persistent and semi-persistent plant viruses, persistently transmitted plant viruses become internalized (enter tissue systems), thus the virus must overcome multiple insect membrane barriers and ultimately infect the salivary gland system for plant inoculation to occur during feeding [3, 4]. Insect gut is the first major barrier that these persistent viruses have to encounter during their entry into the vectors [3, 5]. Despite the fact that many viral determinants of transmission have been characterized [6], the key molecules of vectors that govern the initial infection of vector guts are just beginning to be characterized. With the advance of high-throughput sequencing, the global transcriptional responses of several insect vector species to their transmitting plant viruses were investigated [7–16]. However, most advances in these studies were heavily focused on the responses of whole-body insects to virus infections with little work centered on the gut tissue level [9, 10, 17]. Given that the vector competence for many persistent viruses is initially dependent on infection of the gut tissue, understanding the molecular interactions between vector gut and virus becomes essential.

*Tomato spotted wilt virus* (TSWV), type species in the genus *Orthotospovirus* (Family *Tospoviridae*, Order *Bunyavirales*), causes serious diseases in U.S. and global agriculture. TSWV is exclusively transmitted by herbivorous thrips species (Thysanoptera: Thripidae). By virtue of the wide geographical distribution and efficiency in virus transmission, *Frankliniella occidentalis* Pergande (the western flower thrips), is considered as the primary vector of TSWV [4]. TSWV consists of a tripartite genome of negative-sense, single-stranded RNAs (denoted as L, M, and S) that are encased in a lipid bilayer envelope. Each genome segment is encapsidated by viral nucleoprotein (N) and bound to viral RNA-dependent RNA polymerase (L). The viral glycoproteins (Gn and Gc) that decorate the virus particle are the viral determinants of vector acquisition in the midgut [18–21]. Two non-structural proteins, NSm and NSs, are also encoded by TSWV functioning as a movement protein in plant hosts and a silencing suppressor in plant and insect cells, respectively [22–24].

TSWV is a persistent-propagative virus, of which transmission is tightly linked to the development of *F. occidentalis* [15, 25, 26]. TSWV infection can be sustained in the midgut after virus ingestion by adult thrips, but the transmission-competency of thrips presents only if the virus is acquired as larvae [27]. The anterior region of larval midgut is the first site of TSWV infection, where after infection of a few epithelial cells in the first instar larva (L1), TSWV multiplies and disseminates into the adjacent cells and eventually infect the remaining midgut tissue. This results in the continuous increase of virus titer as larva develops into the second instar larva (L2). During *F. occidentalis* development, virus titer reaches its highest peak in the late L2 stage [26]. Following the first cycle of virus accumulation in larval stages, virus titer decreases rapidly in the propupal and pupal stages. The second virus accumulation cycle appears during the adult stage when the principal salivary glands become the primary site for TSWV multiplication [25, 26, 28–30].

The transcriptome responses of whole-body *F. occidentalis* and *F. fusca* to TSWV exposure at various developmental stages (larval, pupal, adult) revealed that TSWV perturbed the transcriptomes of both species in a developmental stage-specific manner. However, the overall responses of both species to the virus exposure were subtle. Less than 1% and 1.6% of transcripts were differentially expressed in *F. occidentalis* and *F. fusca*, respectively [14, 15]. Another proteome study on TSWV-exposed and non-exposed *F. occidentalis* L1s also reported that only 5% of proteins were differentially regulated due to TSWV infection [31]. These findings may imply the nature of non-pathogenicity of TSWV to specific thrips vectors or the presence of potential relationship between viral tissue tropism and the degree of thrips response to TSWV infestation.

Considering the tight linkage between thrips development and TSWV transmission, as well as the determinative role of gut tissue in the initial infection process, we aimed to identify gut genes that are differentially regulated in the course of TSWV infection during larval development. Therefore, we conducted a comparative transcriptome analysis on gut samples of *F. occidentalis* associated with TSWV infection at the three different time points during the larval developmental stages (L1, early L2, and late L2) using RNA sequencing (RNA-Seq). The three time points represent not only the different larval developmental stages, but also the varying levels of virus accumulation and dissemination in the larval guts from a few infected epithelial midgut cells to the general infection of the entire gut [25, 32, 33]. Additionally, we performed a weighted gene correlation network analysis (WGCNA) to identify clusters of co-expressed gut genes (modules) significantly correlated with thrips larval stages and also the potential hub genes responsive to virus infection in the most significant gene modules. Lastly, to gain a deeper insight into the effect of virus on gut gene expression before and after removal of virus sources, a time-course experiment was performed to detect the temporal expression of network hub and peripheral transcripts. To our knowledge, this is the first gut tissue-specific omics study for any thysanopteran species. This work will provide insights into the important genetic components that respond to TSWV exposure in *F. occidentalis* larval guts and also further our understanding of the roles of guts in the thrips- TSWV interactions.

## RESULTS

### Sequence read quality and summary statistics

Over 1.8 billion raw reads in total were generated for the 24 RNA-Seq libraries, and after quality filtering, 80% of the reads on average were retained with 88% of them having Phred quality scores ≥ 30 and an average length of 143bp (**Table S1**). On average across the 24 libraries, 86% of the quality paired end reads aligned to the annotated gene models in the *F. occidentalis* genome reference.

**Table 1.** Top ten most differentially-responsive gut transcripts of tomato spotted wilt virus- infected *Frankliniella occidentalis* larvae

| <sup>a</sup>Time-larval stage | Transcript ID | <sup>b</sup>Log<sub>2</sub> fold change | <sup>c</sup>Adj. p-value | <sup>d</sup>Gene description |
| --- | --- | --- | --- | --- |
| 3-L1 | FOCC009344 | 7.36 | 5E-05 | cuticle protein CP14.6 |
|  | FOCC001013 | 7.29 | 2E-03 | zinc finger protein 512B-like |
|  | FOCC001012 | 6.56 | 1E-02 | hypothetical protein FOCC_FOCC001012 |
|  | FOCC013209 | 6.54 | 7E-04 | low-density lipoprotein receptor-related protein 12 |
|  | FOCC006427 | 6.11 | 1E-05 | golgin subfamily A member 6-like protein 22 |
|  | FOCC001001 | 5.29 | 9E-04 | hypothetical protein FOCC_FOCC001001, partial |
|  | FOCC017932 | 5.17 | 6E-04 | hypothetical protein FOCC_FOCC017932, partial |
|  | FOCC005944 | 4.78 | 4E-03 | proline-rich extensin-like protein EPR1 |
|  | FOCC001010 | 4.73 | 1E-02 | hypothetical protein FOCC_FOCC001010 |
|  | FOCC002062 | -5.02 | 0E+00 | venom carboxylesterase-6-like |
| 24-L2 | FOCC012858 | -1.39 | 2E-05 | myrosinase 1-like |
|  | FOCC015174 | -1.96 | 1E-02 | transcriptional regulator ATRX-like isoform X1 |
|  | FOCC015182 | -2.13 | 3E-03 | hypothetical protein FOCC_FOCC015182 |
|  | FOCC015649 | -2.51 | 3E-02 | hypothetical protein FOCC_FOCC015649 |
|  | FOCC015178 | -2.75 | 4E-03 | death-associated inhibitor of apoptosis 1-like isoform X2 |
|  | FOCC012492 | -4.43 | 5E-02 | hypothetical protein FOCC_FOCC013062 |
|  | FOCC004067 | -7.40 | 2E-02 | foot protein 1 variant 2 |
| 48-L2 | FOCC001020 | -4.72 | 3E-06 | skin secretory protein xP2-like |
|  | FOCC014805 | -4.98 | 5E-06 | endocuticle structural glycoprotein SgAbd-8-like |
|  | FOCC007139 | -5.06 | 3E-03 | glycine-rich cell wall structural protein-like |
|  | FOCC017317 | -5.16 | 5E-06 | endocuticle structural glycoprotein SgAbd-8-like |
|  | FOCC016782 | -5.23 | 3E-02 | farnesyl pyrophosphate synthase |
|  | FOCC001002 | -5.38 | 7E-07 | hypothetical protein FOCC_FOCC001002 |
|  | FOCC014804 | -5.59 | 5E-09 | paternally-expressed gene 3 protein-like |
|  | FOCC016120 | -5.71 | 2E-04 | cuticle protein 7 |
|  | FOCC017781 | -6.80 | 3E-04 | terminal uridylyltransferase 4-like |
|  | FOCC017730 | -9.20 | 3E-02 | hypothetical protein FOCC_FOCC017730, partial |

<sup>a</sup>Time-larval stage = cohorts of first instars (L1) given a 24-hour acquisition access period on virus-infected plant tissue (or healthy tissue for control group), then moved to healthy green bean pods as food source, followed by sampling of larval groups at 3 hours (3-L1), 24 hours (24-L2, early second instar), and 48 hours (48-L2, late second instars) after removal of virus inoculum. <sup>b</sup>RNA-Seq, differential expression analysis of normalized read counts of tomato spotted wilt virus-infected thrips relative to non-infected thrips (sign/no sign indicates direction of change); <sup>c</sup>False discovery rate (FDR) corrected p-values with the Benjamini and Hochberg method [34]; <sup>d</sup>Blastx query of FOCC transcript against NCBI non-redundant protein database using Blast2GO. Refer to **Table S2** for a list of all significant ( $P_{\text{adj}} < 0.05$ ) differentially-expressed transcripts and annotations.

### Differential gene expression of larval guts in response to TSWV

There were 147 non-redundant DETs in response to TSWV over the course of larval growth and development, of which 65% were associated with L1 (**Figure 1**). The number of up and down- regulated DETs in L1 guts was comparable; however for both of the L2 stage-times, most to all of the DETs were down-regulated relative to their non-infected counterparts (**Figure 1A**). The least perturbed stage-time was early L2s, 24 hours after removal of inoculum, with only 7 DETs. Across the three stage-times, the absolute change in transcript expression between virus-infected and non-infected ranged from 2 to 588 folds (1.04 – 9.2 log2fold), and the average log2fold change ranged from -3.68 (48-L2) to 3.41 (3-L1). The top 10 most virus-responsive transcripts and their predicted annotations for each stage-time are listed in **Table 1**.

**Figure 1.**
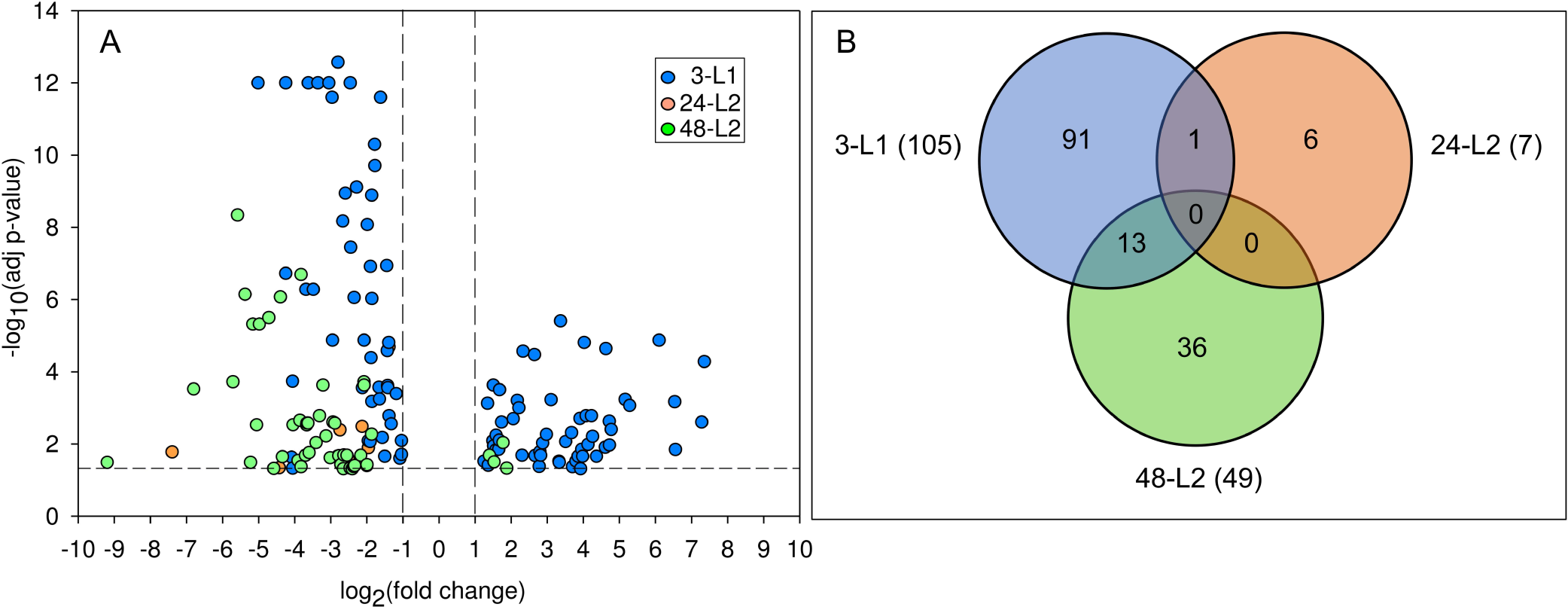
Differentially-expressed transcripts in *Frankliniella occidentalis* larval guts in response to tomato spotted wilt virus infection. (A) Volcano plot showing the distribution of differentially- expressed transcripts (DETs) in virus-infected guts relative to non-infected guts at three different times over larval development: 3-L1 = cohort of first instars three hours after removal of virus inoculum after a 24-hour acquisition access period; 24-L2 = same cohort of insects developed into early second instars, 24 hours after removal of inoculum; and 48-L2 = same cohort of insects entering into late second instars, 48 hours after removal of inoculum. The horizontal and vertical dotted lines indicate cutoffs for log2-fold change ≥1 and FDR adjusted *P*-values ≤ 0.05, respectively. (B) Venn diagram depicting the number of shared and unique DETs between the three time-larval stages; numbers in parentheses indicate number of DETs in each time-larval stage.

Of the 147 DETs, 18.4% of the sequences matched hypothetical proteins (**Table S2**), whereby the most significant matches were NCBI Genbank accessions of the *F. occidentalis* OGS v1.0 peptides, as well as hypothetical proteins of *Thrips palmi*, another genome-sequenced thysanopteran. In addition, only 3.4% of the DETs matched uncharacterized proteins in other metazoans. Only four DETs in the total dataset did not contain predicted Interpro IDs specifying conserved peptide motifs, e.g., transmembrane domain, cytoplasmic domain, coil, and these sequences belonged to the ‘hypothetical proteins’ set.

**Table 2.** Intramodular network hub gene connectivity in co-expression modules enriched with differentially-expressed transcripts responsive to tomato spotted wilt virus.

| Module network | Hub transcript ID | <sup>a</sup>kME; kIN | Gene description | <sup>b</sup>Log<sub>2</sub> fold change |  |
| --- | --- | --- | --- | --- | --- |
|  |  |  |  | 3-L1 | 48-L2 |
| Turquoise | FOCC013756 | 0.801; 8.036 | cytochrome p450 (<sup>c</sup>CYP3653A2) | -1.09 | - |
|  | FOCC012444 | 0.685; 6.274 | uncharacterized protein | -1.33 | - |
|  | FOCC012265 | 0.693; 5.767 | myrosinase 1-like | -1.65 | 1.88 |
| Blue | FOCC009188 | 0.969; 9.711 | hypothetical protein | 1.64 | - |
|  | FOCC004976 | 0.938; 9.121 | lysozyme X-like isoform X1 | 1.65 | - |
|  | FOCC013675 | 0.885; 6.677 | lysosomal alpha-mannosidase | 1.25 | - |
| Light cyan | FOCC014803 | 0.981; 17.448 | <sup>d</sup>endocuticle structural glycoprotein | 3.33 | -3.83 |
|  | FOCC017589 | 0.980; 16.886 | cytochrome p450 (<sup>c</sup>CYP4G236) | 2.82 | - |
|  | FOCC007741 | 0.985; 17.065 | troponin C, isoform 1 | - | -2.89 |
|  | FOCC001084 | 0.977; 16.152 | cuticle protein 6-like | - | -3.69 |

<sup>a</sup>module membership (kME) and intramodular connectivity (kIN) calculated using methodology from Langfelder and Horvath, 2008 [36] (WGCNA R package, updated version February 16, 2016); <sup>b</sup>RNAseq analysis of normalized read counts between tomato spotted wilt virus-infected thrips relative to non-infected thrips; <sup>c</sup>CYP nomenclature from Rotenberg et al., 2020 [35]; <sup>d</sup>TSWV-interacting protein, endoCP-G<sub>N</sub> [33] .

### Developmental stage-dependent larval gut response to TSWV

There was little to no overlap in virus-responsive DETs across the three larval stage-times (**Figure 1B**). The two stages farthest apart in developmental time (3-L1 and 48-L2) shared the most in number of DETs (13), which might be expected given the few number of DETs associated with the early L2 stage (24-L2). The 13 shared DETs have provisional gene ontologies of proteolysis and hydrolysis, carbohydrate metabolism, lipid metabolism, and structural constituent of cuticles, such as transmembrane protease serine, putative beta-glucosidase, myrosinase, acyl-CoA desaturase, and endocuticle structural glycoprotein. Most of these 13 transcripts were down- regulated in both larval stage-times. Of those 13, five disagreed in direction between 3-L1 and 48- L2, and these were annotated to be involved in lipid metabolism and structural constituent of cuticle. The single DET shared between 3-L1 and 24-L2 was putatively predicted to be a myrosinase, down-regulated in both stages.

### Gene ontologies of DETs mirror developmental stage-dependent response to virus

Of the 147 DETs in larval guts, 58% were significantly assigned GO terms. These GO-annotated sequences were classified by biological process (BP, 64 sequences), molecular function (MF, 81 sequences), and cellular localization (CC, 17 sequences) (**Figure 2, Table S1**). Reflecting the little overlap of DETs between the developmental stages, enriched GO terms associated with these DETs also varied across stages. Distributed across the three types of GO classifications, the most commonly occurring annotations for guts of the 3-L1 stage were ‘oxidation reduction process’, ‘carbohydrate metabolic process’, ‘hydrolase activity’, and ‘integral component of membrane’, while the predominant GOs in the 48-L2 stage indicated proteins associated with ‘lipid metabolic process’, ‘structural constituent of cuticle’, and ‘extracellular region’ and ‘integral component of membrane’. Proteins with GO annotations of ‘transport’ and ‘heme-binding’, and ‘iron-binding’ were identified only for 3-L1 guts. The DETs associated with roles in ‘transport’ included transmembrane proteins that transport sugars (Tret1-2) and solutes (MFS-type transporter) or cations (organic cation transporter), and proteins that transport lipids in the hemolymph to various tissues (apolipophorins). The ‘heme-binding’ and similarly the ‘iron-binding’ functional group was comprised of cytochrome p450s (CYP gene superfamily) belonging to clans 3 (typically CYP6) and 4 (typically CYP4), including two sequences (FOCC013756 and FOCC015103) belonging to two newly discovered and curated CYP gene families (CYP3653A2 and CYP3654A1, respectively) from the *F. occidentalis* genome [35].

**Figure 2.**
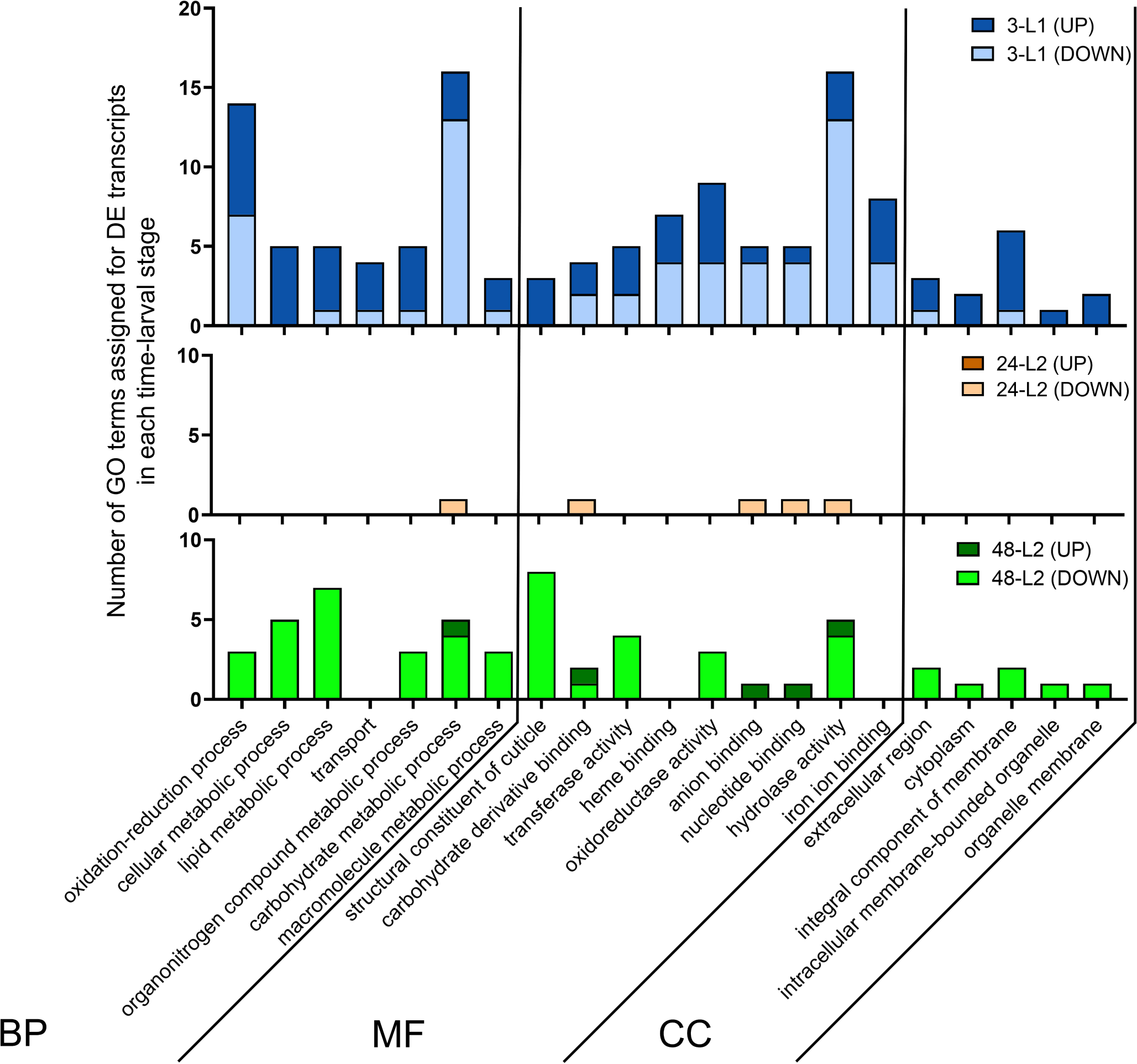
Distribution of up- and down-regulated, tomato spotted wilt virus-responsive gut transcripts of *Frankliniella occidentalis* larvae into provisional gene ontologies (GO) by multi- level GO classification of sequences into biological processes (BP), molecular functions (MF), and cellular component (CC). 3-L1 = cohort of first instars three hours after removal of virus inoculum after a 24-hour acquisition access period; 24-L2 = same cohort of insects developed into early second instars, 24 hours after removal of inoculum; and 48-L2 = same cohort of insects entering into late second instars, 48 hours after removal of inoculum.

### Validation of the RNA-Seq analysis

Using dissected 3-L1 guts from a similarly conducted and replicated TSWV acquisition assay, it was determined that 80% of a subset of 10 selected DETs were validated with ddPCR in direction and, for the most part, the magnitude of change in expression as compared to RNA-Seq (**Figure 3**). A correlation analysis of the log2fold-change values yielded a significant Pearson correlation coefficient of 0.72 (*P* = 0.0185), indication of significant validation of the gut RNA-Seq.

**Figure 3.**
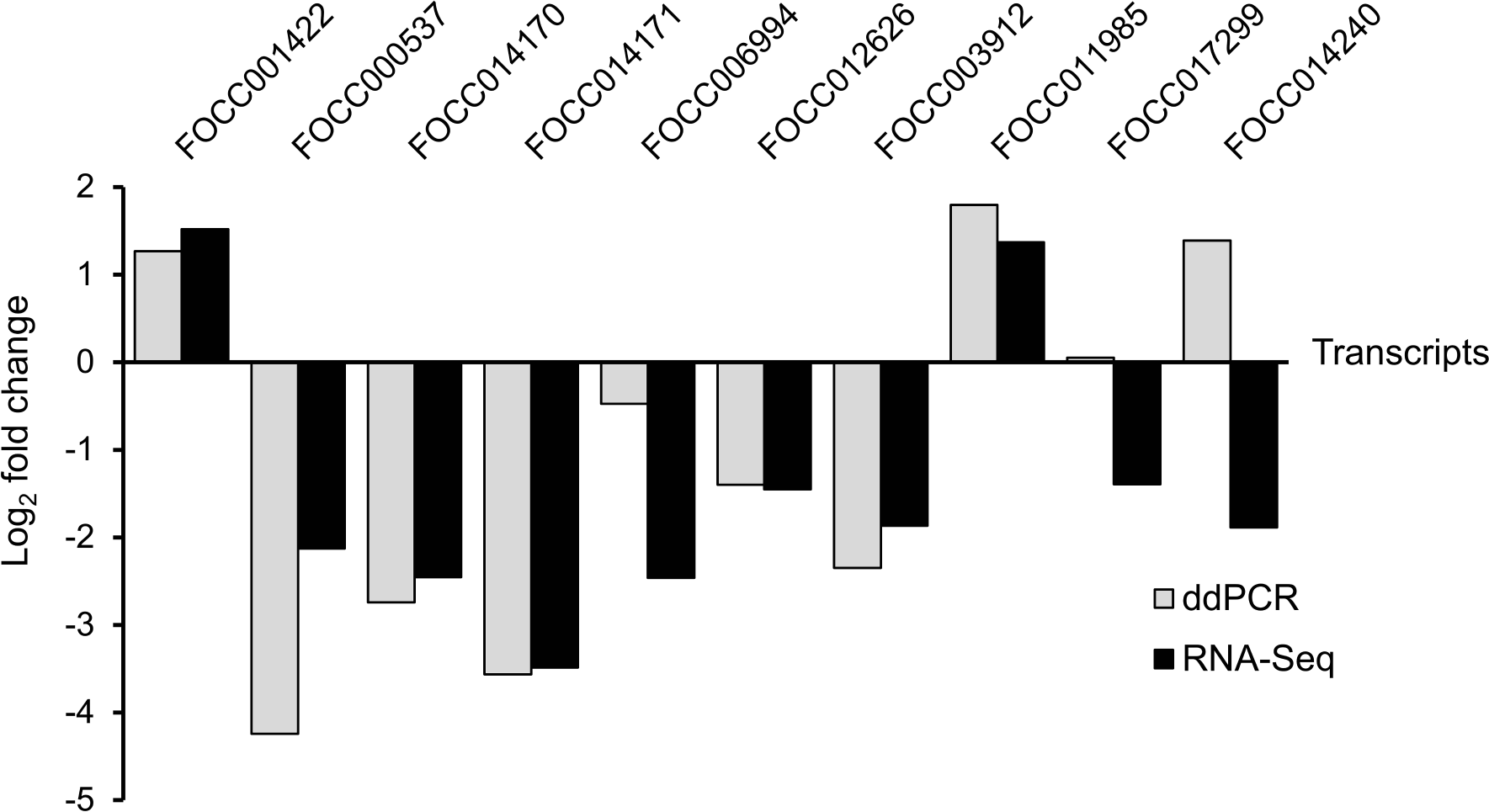
Digital-droplet PCR (ddPCR) validation of the RNA-seq analysis using selected differentially-expressed transcripts (FOCC prefix) of *Frankliniella occidentalis* first instar larval (L1) guts in response to tomato spotted wilt virus infection. Each transcript was normalized to *F. occidentalis* actin RNA and calibrated to non-infected L1 guts; each bar represents a pool of 100 L1 guts sampled after a 24-hour acquisition access period followed by a 3h-clearing on green bean pods.

### Network and intramodular membership analysis of co-expressed DETs revealed highly interconnected hub genes

WGCNA of normalized read count data across the larval gut transcriptome clustered co-expressed transcripts into 31 discrete modules (**Figure 4A**, module eigengenes) that formed four main clades (**Figure 4B**). Of these modules, 5, 3, and 4 were significantly correlated (*P* < 0.05) with the 3-L1, 24-L2, and 48-L2 stages, respectively (**Figure 4A**, absolute values of the correlation coefficient). The larval gut genes responsive to virus infection (147 DETs) were distributed across 14 modules (**Figure 4C, Table S3**), and collectively held intramodular membership across the four main clades (**Figure 4B**). The DETs nested within each time-stage were distributed across 10, 3 and 10 shared and non-overlapping modules for 3-L1, 24-L2 and 48-L2, respectively. The most DET-enriched modules were the ‘lightcyan’ (44 sequences), ‘turquois’ (41 sequences), and ‘blue’ modules (31 sequences) (**Figure 4C**), together accounting for 78.9% of the gut DETs. For the most part, the ‘turquoise’ module DET members were down-regulated, regardless of stage, and both the ‘blue’ and ‘lightcyan’ modules consisted of DET members that were split in direction of change along the lines of developmental stage, with most 3-L1 DETs up-regulated and all L2 DETs down- regulated in these modules. The three enriched modules resided in different main clades (**Figure 4B**), indication of distant relationships or very little biological connection between these modules [36]. The ‘turquoise’ and ‘blue’ modules were predominantly occupied by 3-L1-associated DETs (83.7% and 74.2%, respectively), while the ‘lightcyan’ was occupied comparably by DETs associated with 3-L1 and 48-L2 (56.3% and 43.8%, respectively) (**Figure 4C, Table S3**). With only 7 DETs associated with 24-L2, most of them were associated with the ‘blue’ module.

**Figure 4.**
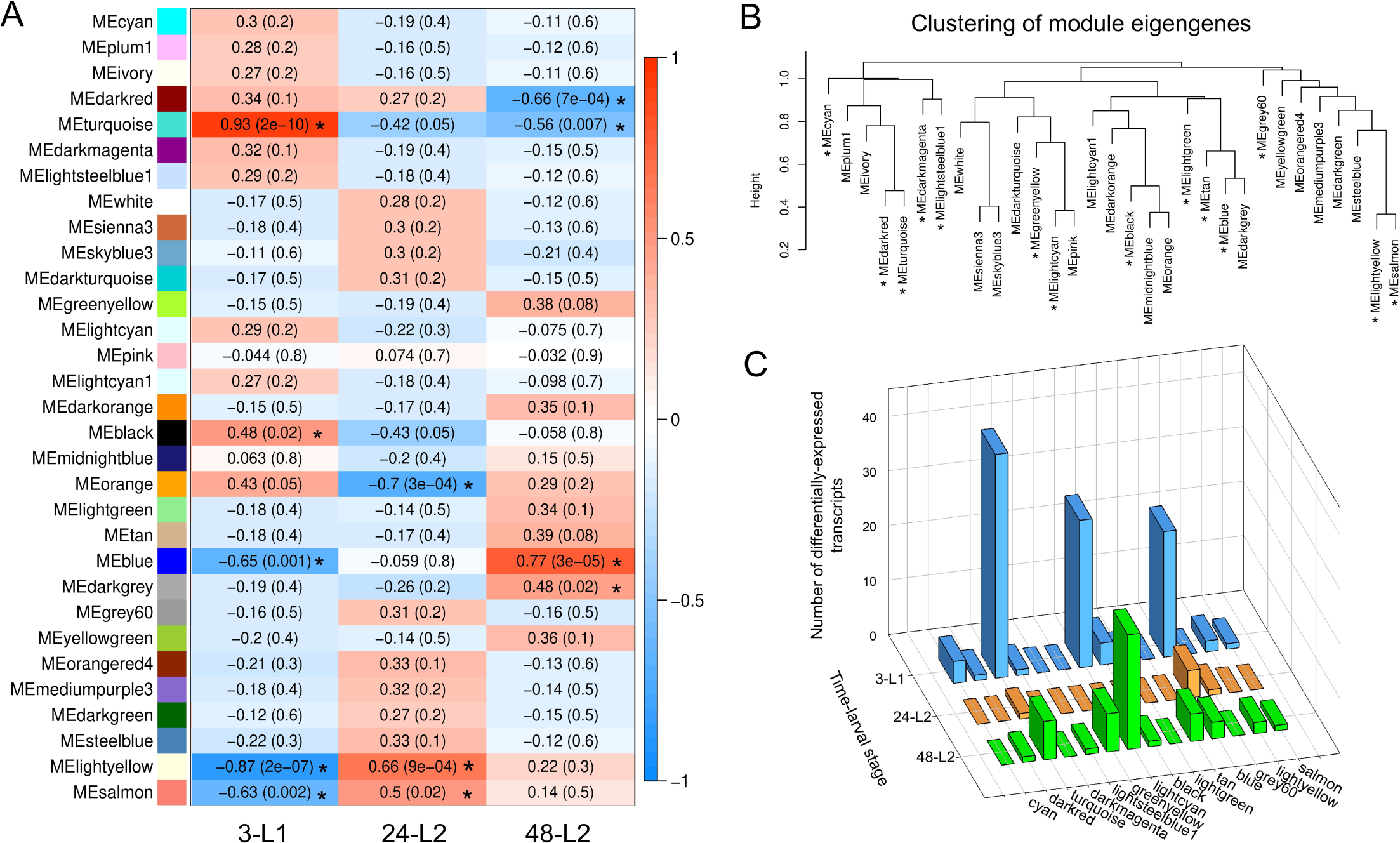
Weighted gene co-expression network analysis (WGCNA) of gut-expressed transcripts in *Frankliniella occidentalis* to identify modules of co-expressed transcripts associated with larval development and response to tomato spotted wilt virus infection. (A) Module-trait relationships depicting significant associations (correlations, *P* < 0.05) between clusters (modules) of co- expressed transcripts (left-hand column of colored blocks, i.e., module eigengenes) and time-larval stages (trait). Each trait includes normalized read counts of all gut-expressed transcripts from four biological replications and two conditions (both virus-exposed and non-exposed) of the gut RNA- seq analysis, i.e., eight samples per time-stage. Asterisks indicate modules that are significantly associated with time-larval stages. (B) Hierarchical clustering of module eigengenes. The dendrogram depicts relationships among modules (eigengenes) of co-expressed transcripts; positively correlated eigengenes cluster within the same clade. Asterisks indicate modules that include sets of differentially-expressed transcripts (DETs) in response to tomato spotted wilt virus infection. (C) Counts of larval gut DETs across 14 modules show enrichment in the turquois, lightcyan and blue modules. 3-L1 = cohort of first instars three hours after removal of virus inoculum after a 24-hour acquisition access period; 24-L2 = same cohort of insects developed into early second instars, 24 hours after removal of inoculum; and 48-L2 = same cohort of insects entering into late second instars, 48 hours after removal of inoculum.

Examination of gene descriptions and GO annotations revealed that the turquoise module represents proteins involved in breakdown of harmful exogenous compounds (e.g. toxic plant substances and insecticides), thermal or antiviral response, proteolysis, and hydrolysis of plant-derived secondary metabolites, collectively considered to describe detoxification and host defense activities. The ‘blue’ module contains proteins known in other insects to be involved in protein, lipid and sugar transport and storage in preparation of quiescent periods or response to environmental stress, and proteolysis and deglycosylation of proteins in the lysosome – collectively considered to describe metabolic energy economics, *i.e.*, reserves and recycling. The ‘lightcyan’ module consists of proteins associated with cuticles (many cuticle proteins and endocuticle glycoproteins) and membranes, lipid metabolism, several transcription factors, and neuropeptides, collectively considered to be consistent with insect growth and development.

Three DET hubs were identified in each of the ‘turquoise’ and ‘blue’ modules and four were identified for the ‘lightcyan’ module (**Figure 5, Table 2**). Module memberships, *i.e.*, the correlation of a node to the module eigengene, of these 10 hubs ranged from 0.685 – 0.985 depending on the module (**Table 2**). The DET hubs with the highest connectivity within their modules were CYP3653A2 (FOCC013756) in the ‘turquoise’ module, a hypothetical protein (FOCC009188) in the ‘blue’ module, and an endocuticle structural protein - specifically endoCP- GN (FOCC014803), a TSWV-interacting proteins identified in *F. occidentalis* L1s [33] - in the ‘lightcyan’ module (**Table 2**, **Table S4**). The ‘turquoise’ and ‘blue’ DET hubs exhibited subtle change due to virus infection (∼2- to 3-fold change), while the four ‘lightcyan’ hubs exhibited a more pronounce effect in response to virus regardless of developmental stage (∼7- to 14-fold change) (**Table 2**).

**Figure 5.**
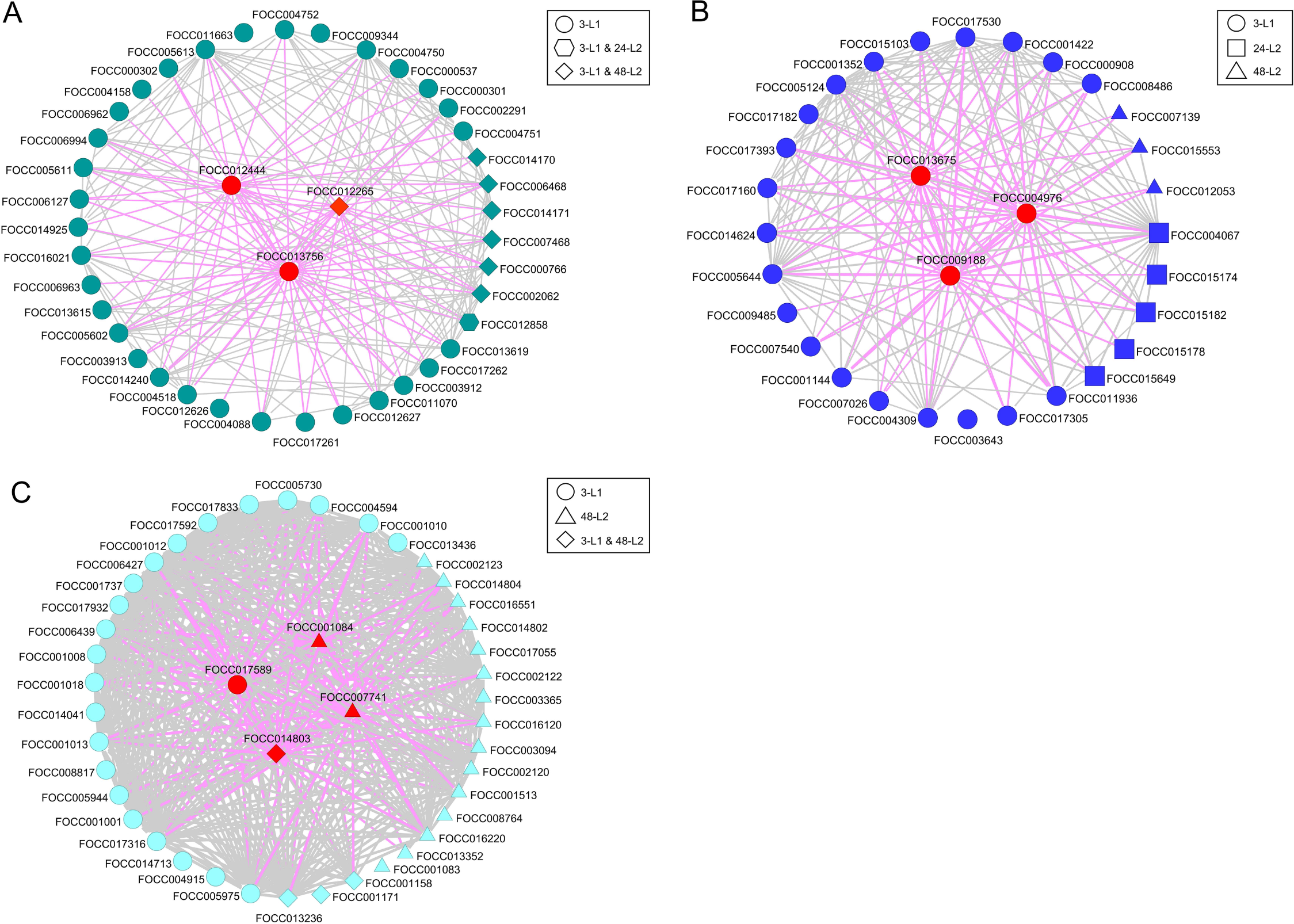
Three intramodular gene networks of differentially-expressed transcripts (DETs) of *Frankliniella occidentalis* larval guts in response to tomato spotted wilt virus infection during growth and development. Together, the turquoise (A), blue (B) and lightcyan (C) modules accounted for ∼80% of the DETs associated with virus infection. DETs associated with time-larval stages (3-L1, 24-L2, and 48-L2) are indicated. Transcripts that were in the top 10% of all module transcripts with regards to both module membership (kME, correlation of a node to a module eigengene) and intramodular connectivity (kIN, sum of connection weights between a node and all its network neighbors) were considered candidate hub transcripts (red shapes). Grey and pink lines indicate the correlations between transcripts and interconnections between transcripts and hubs, respectively. For better visualization of the interactions, a weight cutoff of ≥ 0.15 was applied for turquoise and blue networks, and a weight cutoff of ≥ 0.35 was applied for the light cyan network. Line thickness indicates strength of the connection. 3-L1 = cohort of first instars three hours after removal of virus inoculum after a 24-hour acquisition access period; 24-L2 = same cohort of insects developed into early second instars, 24 hours after removal of inoculum; and 48-L2 = same cohort of insects entering into late second instars, 48 hours after removal of inoculum.

Notably, there were a few tightly connected DETs assigned to other modules (**Table S2**). For instance, the FOCC006197 and FOCC000732 assigned to the ’cyan’ and ’greenyellow’ modules, respectively, showed the highest module membership in each module, and the second highest connectivity among the 144 transcripts in ’cyan’ module and 191 transcripts in ’greenyellow’ module (**Table S2**, “cyan” and “greenyellow” module tabs). Moreover, after a weight cutoff of 0.15, both transcripts established the highest number of connections (89 and 119 for FOCC006197 and FOCC000732, respectively) with other module members, which altogether indicates that they likely play an important role in influencing the expression of co-expressed genes and the subsequent biological processes.

### Significant interactions revealed between time and virus treatment on larval gut expression for selected network hub and connecting DETs

Gut expression patterns were documented for developing larvae during and after removal of the virus inoculum for three of the four network DETs examined (**Figure 6A, C, D**). Coinciding with the shift in larval stage (molting) from L1 to L2, a significant shift in expression pattern of non-infected guts was observed between 30h and 42h for FOCC013675 (blue hub: lysosomal alpha-mannosidase, *P* = 0.0004), FOCC013756 (turquoise hub: cytochrome P450, CYP3653A2, *P* < 0.0001), and FOCC001422 (blue network DET: putative beta-glucosidase 6, *P* = 0.0003). The shifts in expression patterns of virus-infected guts were not significant for any of these DETs. Additionally, there was a significant treatment*time interaction for FOCC013675 (*P* = 0.0005), FOCC013756 (*P* < 0.0001), and FOCC001422 (*P* = 0.0004), indicating that the effect of virus treatment depended on time of sampling, and incidentally, larval age and stage. Indeed, the effect of virus on gene expression reversed in direction between 30 and 48 hours after L1s first encountered virus-infected tissue, which was equivalent to 6 (early L2) and 24 hours (mid L2) after removal of the inoculum. Consistent between the three temporally-dynamic DETs, the virus effect first occurred between 6 and 18 hours during exposure to virus-infected tissue, and the direction of change in expression confirmed findings obtained for the RNA-Seq time-stages. The virus effect was documented during feeding on infected tissue and after removal of the inoculum (*P* < 0.05; **Figure 6 A, C, D**), however, with some variation depending on the DET. For example, transcript abundance of CYP3653A2 was significantly modified by virus at 5 of the 6 sampling time points, while lysosomal alpha-mannosidase and the beta-glucoside transcripts exhibited response to virus at 3 time points at varying times. For the fourth DET monitored - lysozyme (FOCC004976), there was no significant change in expression over time (*P* = 0.5186) nor a treatment*time interaction (*P*=0.6756), and the amount of variation resulted in no apparent virus effect. However, it was more stably expressed in virus-infected guts compared to non-infected control, indicated by their relatively less variation in expression (**Figure 6B**).

**Figure 6.**
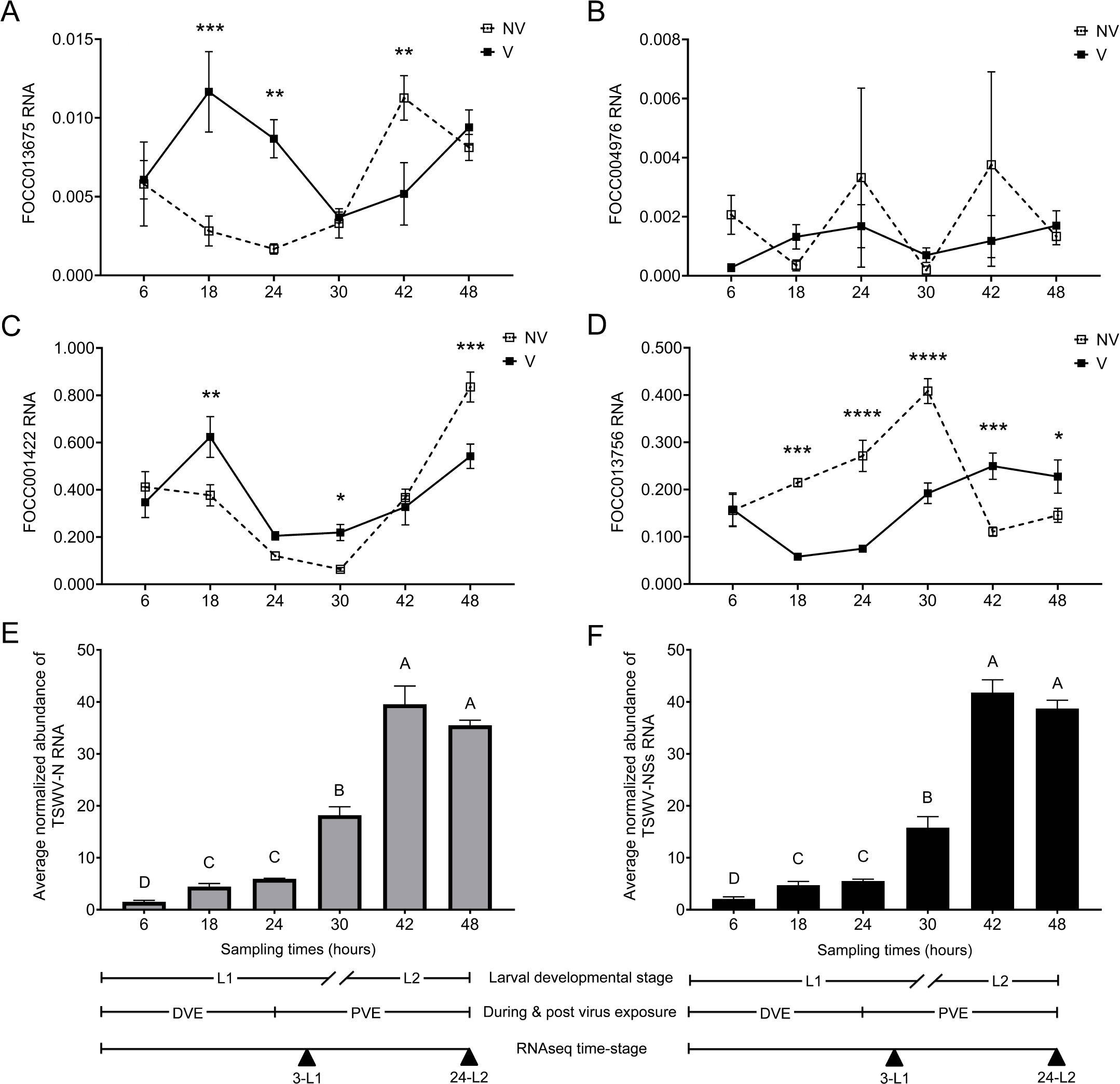
Temporal gut-expression of selected hub and connecting transcripts in the blue and turquoise co-expression gene networks for non-exposed (NV) and tomato spotted wilt virus (TSWV) - infected (V) larvae of *Frankliniella occidentalis*. Digital-droplet PCR was performed to quantify normalized transcript abundance of A) FOCC013675 (lysosomal alpha-mannosidase, blue hub), B) FOCC004976 (lysozyme, blue hub), C) FOCC013756 (cytochrome p450: CYP3653A2, turquoise hub), and D) FOCC001422 (putative beta-glucosidase, blue connecting gene) during (DVE) and post virus exposure (PVE) at six time points during larval growth and development. Asterisks indicate statistically significant differences in gene expression level for pairwise comparisons between V and NV guts (\**P* < 0.05, \*\**P* < 0.01, \*\*\**P* < 0.001, \*\*\*\**P* < 0.0001). The corresponding abundance of E) TSWV nucleocapsid gene (N RNA) and F) TSWV non-structural small RNA gene (NSs RNA) in larval guts was also temporally monitored using real-time quantitative reverse-transcription PCR (qRT-PCR). Normalized abundance of each thrips and viral RNA was normalized to *Frankliniella occidentalis* actin RNA. Bars headed by different letters indicate statistically significant differences (*P* < 0.05) between time-points. No virus was detected in NV guts. Developmental stage (L1, L2) and time-stage corresponding to the RNAseq analysis of this study are delineated at the bottom of the figure; double dashes in the developmental stage timeline indicate the transition period (molting) from L1 to L2, thus the 30- hour sample included both stages. Each expression data point/bar represents the mean and standard error of three biological replicates (independent experiments), and each replicate represents a pool of 60 larval guts.

Virus accumulation, approximated by either normalized abundance of TSWV N (**Figure 6E**) or TSWV NSs (**Figure 6F**), was also documented in larval guts during exposure and after removal of inoculum. TSWV RNA significantly increased 3.3-fold in average over the course of feeding on infected tissue (L1-6h and L1-24h comparison, *P*<0.0001 for N and *P*=0.0014 for NSs) and 2.2-fold after removal of the inoculum (L1-24h and L2-24h comparison, *P*=0.0093 for N and *P*=0.0035 for NSs). A correlation analysis of TSWV RNA abundance and DET expression showed a significant correlation between virus abundance and CYP3653A2 gene expression with a correlation coefficient of 0.5968 between N and CYP3653A2 (*P*=0.0089), and 0.6311 between NSs and CYP3653A2 (*P*=0.005). There were no significant correlations between virus abundance and the other DETs.

## Discussion

The insect alimentary canal is a structurally complex organ that plays a vital role in food digestion and absorption, and in promoting or suppressing microbial and viral associations [37–39]. Previous -omics studies on thrips vector species in response to orthotospovirus infection have primarily focused on the whole-body level [13–15, 31], with a common finding that transcriptome-level responses vary greatly over the course of thrips development from larvae to adulthood. In this study, we similarly found a tight link between developmental stages (L1 and L2) and gut transcriptome responsive to TSWV, which may be due to gut activities and physiological changes that occur between the two larval stages (L1 and L2), reflected by the number, diversity and putative functional roles of the differentially-expressed transcripts between the larval stages.

Consistent with the previous whole-body transcriptome studies on *F. occidentalis* and *F. fusca* [14, 15], the responses of *F. occidentalis* larval guts to TSWV infection were subtle (less than 1%) in terms of the number of significantly altered gene expressions. However, the magnitude of responses (fold change in expression) in larval guts was significantly increased by 7.5-fold in average compared to whole-body *F. occidentalis* larvae (*P*<0.0001, Mann-Whitney test). This suggests the presence of a dilution effect when pooling all thrips tissues where viruses are only limited to certain tissues/cells, emphasizing the potential linkage between viral tissue tropism and thrips response to virus infestation.

Insects have evolved diverse defensive mechanisms to cope with the accumulation of harmful substances in their tissues [40]. Comparison of the provisional functions of all DETs across the larval stages revealed a distinct difference between L1 and L2 guts, where three GO annotation groups were uniquely identified in L1 gut but not in L2 (**Figure 2**). For example, the organic cation transporter assigned to the GO annotations of ‘transport’ plays an important physiological role in secretion of organic cations derived from endogenous and exogenous origin, including toxic environmental compounds, pharmacological drugs, and plant defense chemicals [41]. Mutation of an organic cation transporter gene in *Drosophila melanogaster* was also shown to cause apoptotic cell death [42]. Cytochrome P450s (P450s) have long been considered multi- functional roles in the oxidative metabolism of a wide variety of plant secondary metabolites and pesticides, as well as in biosynthetic pathways of juvenile hormone and ecdysteroids, which are associated with insect growth, development, and reproduction [43–45]. We found that all P450- coding DETs assigned to the GO annotations of ‘heme-binding’ and ‘iron-binding’ were identified in 3-L1 guts but not in L2 guts. Among them, two were newly discovered to be thrips-specific [35], one of which was further predicted to be a ‘turquoise’ hub transcript in our network analysis (**Figure 5A**). Another set of DETs that are involved in detoxification is glutathione S-transferase (GST), which is known to detoxify the xenobiotics by catalyzing the conjugation of glutathione with electrophilic substrates [46]. These GSTs were also exclusively identified in 3-L1 guts. Notably, the perturbed transcripts (down-regulation) coding for these detoxification proteins are only detected in L1 guts shortly after exposing to virus inoculum but not after 24h and 48h post exposure to inoculum, which may indicate the indirect plant-mediated effect of virus on the larval gut, or alternatively their potential involvement in advancing virus infection of the gut or defending against the infection. Further functional studies are required to elucidate their exact roles in thrips and in virus infection of vectors, in particular the thrips-specific, P450 hub gene.

Pathogen invasion into insects stimulates a complex immune response [47]. Understanding the defense response of insect vectors to virus infection is important to uncover the molecular mechanism by which virus overcomes the host immune system. Our study found several immune-related DETs that are up-regulated in virus-exposed guts, including apolipophorins, lysozyme, and trypsin, but only during the early stage (3-L1) of the infection. In addition to their primary role in lipid transport, apolipophorins also function in pattern recognition and innate immune response by stimulating anti-bacterial/fungal activities, phagocytosis, superoxide production, and detoxification of endotoxins of the insect pathogenic bacteria [48–51]. Lysozyme possesses ubiquitous antibacterial enzyme activity [52, 53]. Trypsin is known for its proteolytic activity in food digestion and degradation of pathogens as a defensive response [54]. On the contrary, it was also shown to promote the virus infection in the insect midgut. For instance, the affinity of La Crosse virus (*Bunyavirales,* the same order that includes TSWV) to *Ochlerotatus triseriatus* midgut cells was shown to be increased after *in vitro* proteolytic processing of the virion surface [55, 56]. Similarly, midgut trypsin activity was demonstrated to be required for achieving optimal infection of dengue virus-2 (family *Flaviviridae*) in *Aedes aegypti* [57]. Furthermore, as the virus accumulated in the larval gut of the present study, we observed that some immune-related DETs were down-regulated in the L2 stage. In the early L2 stage (24-L2), an inhibitor of apoptosis (the death-associated inhibitor of apoptosis) was identified to be down-regulated by almost 7-fold in virus-infected guts. Apoptosis is a highly regulated process, and viral infection could induce the programmed cell death response in host cells as a means of eliminating viral pathogens. On the other hand, some viruses may utilize such apoptotic cell death mechanisms to kill cells for their dissemination [58, 59]. In both cases, viruses must block or delay cell death until neighboring cells are invaded. At a time when TSWV abundance has not peaked (as is the case in early L2s), inhibition of apoptosis in thrips gut cells may benefit viral replication to accumulate sufficient progeny for cell-to-cell spread. This may also support the notion that TSWV is non-pathogenic to

*F. occidentalis*. Lastly, in the 48-L2 stage, some DETs involved in anti-microbial/fungal/viral response were down-regulated in virus-exposed guts, such as chitinase, diptericin, and troponin. Another study showed that silencing of the diptericin gene regulated by one of the canonical signaling pathways (immune deficiency or IMD) led to the significant increase in Sindbis virus replication in *Drosophila melanogaster* [60]. We speculate that the down-regulation of diptericin in *F. occidentalis* larval guts may contribute to the accumulation of TSWV.

Weighted gene correlation network analysis (WGCNA) has been widely used to facilitate the understanding of various biological processes in different organisms and to identify highly connected hub genes as candidate biomarkers or therapeutic targets of diseases [36]. Our study findings provided several candidate genes for studying the vector competence of *F. occidentalis*. Mirroring our gene ontology study, a great number of DETs encoding detoxification proteins were clustered into the ‘turquoise’ module, and all DETs associated with lipid transport and storage were clustered into the ‘blue’ module. Similarly, DETs encoding cuticle/endocuticle proteins and others involved in lipid metabolism were clustered into the ‘lightcyan’ module (**Table S4**). To some extent, these findings confirm the validity of our network-based systems biology analysis. As the hub genes share the most connections with other co-expressed genes in a module, understanding their functional roles will likely lead to important biological insights. Although it is uncertain how the hub transcripts crosstalk with other clustered DETs, the tight co-regulation and functional relatedness point to their significance in TSWV infection events. The 10 hub transcripts identified through our network analysis (**Table 2, Figure 5**) prioritize future evaluation to decipher their functions in thrips and in virus transmission using functional analysis tools, such as RNA interference and interactive proteomics. Particularly, CYP3653A2 and FOCC009188 encoding a novel P450 and a hypothetical protein are the two most connected hub transcripts in ‘turquoise’ and ‘blue’ module networks, respectively. Both proteins are uniquely identified in *F. occidentalis*, which make them interesting and important targets to investigate in the future in the context of TSWV transmission.

Our network analysis identified a gut DET coding for endocuticle structural glycoprotein (endoCP-GP, FOCC014803) as the most connected hub in the ‘lightcyan’ network (**Figure 5C, Table 2**). A pairwise alignment of coding sequences (CDS) revealed that FOCC014803 shared 99.9% nucleotide sequence identity (100% coverage) with EndoCP-GN (GenBank accession no. MH884757), an endocuticle structural protein reported to be expressed in *F. occidentalis* larval guts, including the midgut, and interacted directly with TSWV glycoprotein (GN), the viral attachment protein [33]. It is curious that an endoCP-GP containing a chitin-binding domain was expressed in the thrips midgut, the site of TSWV entry, since it is understood that the thysanopteran midgut secretes an extra-cellular lipoprotein membrane devoid of chitin, i.e., perimicrovillar membrane (PMM) [61]. The role of TSWV-binding and/or infection-responsive endoCP-GPs expressed in the midgut have yet to be determined. While it is still unknown whether EndoCP-GN serves as a receptor for TSWV, its perturbation in infected L1 guts coupled with high connectivity to other DETs (i.e., hub gene) underscores a critical role during the virus lifecycle.

Both TSWV accumulation and expression of TSWV-responsive gut genes (DETs) varied over larval development. TSWV accumulation in larval guts supported findings reported for larval whole bodies [26], where the abundance of N RNA increased significantly over larval growth and development (**Figure 6E, F**). In the present study, accumulation of NSs RNA provided confirmatory evidence of increased virus abundance over larval development. While there was no consistent association between virus abundance and DET expression in guts, virus infection did temporally modulate DET expression patterns as compared to non-infected guts. This was most evident at or around molting (transition from L1 to L2) (**Figure 6**). While only a few DETs were monitored in the temporal study, consistently the largest differential between TSWV-infected and non-infected guts occurred i) during exposure to the virus inoculum, and ii) when guts supported the highest virus titers, i.e., during the L2 stage, when the insect was far removed from virus-infected plants. Our temporal study was not intended to completely uncouple gut responses due to direct effects of virus infection from plant-mediated, indirect effects of feeding on infected tissue, however it did reveal patterns of expression that may be experienced by emergent L1s feeding on infected plants in natural settings. Findings from our temporal study warrant further investigation to uncouple direct from indirect effects of virus on larval gut responses during thrips development.

## CONCLUSIONS

Our study centers on the first tissue barrier to internalization of orthotospoviruses into the thrips vector body – the larval gut – and documents transcriptome-level gut response to viral invasion and accumulation during larval growth and development. The key determinant of vector competence for thrips vectors is acquisition of virus during the larval stage, a requisite that uniquely distinguishes thrips vectors from other insect vectors (e.g., aphids, their closest relatives). Our study represents the first description and analysis of the transcriptome-wide response of thrips vector gut tissue to an orthotospovirus, and provides promising, highly connected network hub genes for future biochemical pathway-probing of putative biomarkers of vector competence. In addition, we optimized and implemented ddPCR for quantification of thrips gene expression in order to circumvent the challenges of obtaining sufficient amounts of RNA from minute tissue systems dissected from minute insects, and consequently increasing sensitivity over that of qPCR technology. Our findings and *F. occidentalis* larval gut sequences will enable functional genomics studies for better understanding thrips transmission biology and for developing novel ways to disrupt virus transmission.

## Methods

### Thrips colony and virus source of inoculum

The insect colony of *Frankliniella occidentalis* originating from the island of Oahu, HI was maintained on green bean pods (*Phaseolus vulgaris*) in the laboratory following conditions as previously described [62]. TSWV (isolate TSWV-MT2) was maintained on *Emilia sonchifolia* by thrips-transmission and mechanical inoculation, alternately, as described previously [33]. Consecutive mechanical passage of virus was not allowed to ensure the vital transmissibility of virus by thrips. The symptomatic *E. sonchifolia* leaves collected from 12 days post-mechanical inoculation were used for insect acquisition of TSWV.

### TSWV acquisition by larval thrips

To obtain synchronized L1s (0-16 h) for virus acquisition, adult thrips were allowed to oviposit on healthy green bean pods for 24 hours. Bean pods containing thrips eggs were collected by removal of thrips and incubated at room temperature for three days and then bean pods were thoroughly brushed to remove emerged larvae. A subset of synchronized L1s were used for 24h virus acquisition period (AAP) on symptomatic TSWV-infected *E. sonchifolia* leaves, and the remaining L1s were exposed to TSWV-free healthy *E. sonchifolia* leaves for 24h (**Figure S1**). These virus-infected (V) and non-infected (NV) larvae were transferred to two separate containers containing green bean pods. Larvae were allowed to feed and develop on bean pods for 3h, 24h, and 48h, respectively. Virus acquisition and larval development steps were performed at 25-26℃ with a 16h photoperiod.

Virus acquisition efficiency of larvae after 24h AAP was assessed for each biological replicate by testing virus infection status of 10-12 individual thrips larvae, collected from the cohort of virus-infected 24-L2s. RNA was extracted from individual larvae using Chelex 100 (Bio-Rad Laboratories, Hercules, CA) as described by Boonham et al. 2002 [63]. cDNA was synthesized with 11 ul of extracted RNA supernatant using Verso cDNA kit (Thermo Scientific, Wilmington, DE) following the manufacturer’s protocol and diluted into 4-fold with nuclease-free water. The presence of TSWV-N RNA in each sample was detected using the Quantitative Polymerase Chain Reaction (qPCR). Two technical replicates of 20 ul reaction mixture were included in qPCR for each sample, which contained 200 nM of TSWV-N or thrips internal reference Actin gene primers in the SYBR Green Supermix (Bio-Rad, Hercules, CA, USA). Samples with missing and/or high (>35) Ct values were considered as TSWV-negative insects. The primer pairs used in this test were listed in **Table S5**. The reactions were performed on the CFX Connect Real-Time PCR Detection System (Bio-Rad, Hercules, CA) with the two-step amplification protocol followed by a melting cycle. The thermal cycling protocol began with 1 cycle of 95 °C for 30 seconds (s), then 40 cycles of 95 °C for 10 s and 55 °C for 30 s. The melting cycle was processed after the amplification step by raising the temperature from 55 °C to 95 °C by 0.5 °C increase per second. No less than 90% of infection rate was found for TSWV-exposed larvae from each cohort/biological replicate.

### Thrips gut dissection and RNA sample preparation

A subgroup of larvae from each sampling time point was collected into a Petri-dish and placed on ice for anaesthetization. Thrips larvae were decapitated in 60% ethanol on a microslide by cutting between the pro- and meso-thorax segments with a sterile, polymer-coated, #15 scalpel blade (Southmedic, Barrie, ON, Canada). The principal salivary glands, tubular salivary glands, fat body, and other residual parts/tissues were carefully removed from the gut tissue using ultrafine single deer hair (Ted Pella, Redding, CA). Dissected gut tissue, including foregut, midgut and partial hindgut, was lysed immediately in a 200-ul PicoPure extraction buffer (PicoPure DNA isolation kit, Arcturus, Mountain View, CA). 100 guts were pooled per sample and processed for total RNA extraction using a PicoPure RNA isolation kit (Arcturus) as described by the manufacturer. On-column DNase treatment was performed with a RNase-free DNase set (Qiagen, Valenica, CA). The 200-ul PicoPure extraction buffer containing gut RNA was applied onto two PicoPure spin columns and the final eluted RNA samples were combined for maximizing the RNA yield. To minimize any possible batch effects during the experiments, gut dissections were performed under the same laboratory condition during the same time of the day and by alternating 20 virus-exposed larvae and 20 virus-free larvae until achieving 100 guts for both samples. The quantity and quality of RNA samples were evaluated in Agilent 2100 Bioanalyzer system. The average RNA yields ranged from 1655 to 2412 ng with an average RNA Integrity Number (RIN) score of 8.33 (ranged from 7.70 to 8.80), showing no sign of RNA degradation. RNA samples were stored in -80 °C until use. This time-course experiment was repeated four times.

### Library preparation and RNA sequencing

All 24 gut RNA samples were sent to the Genomic Sciences Laboratory facility at North Carolina State University (NCSU) for cDNA library preparation and RNA sequencing. The directional (strand-specific) RNA-seq libraries were constructed with poly(A) mRNA enrichment and rRNA depletion processes using the NEBNext Ultra II Directional RNA Library Prep Kit (New England Biolabs, Ipswich, MA). The indexed libraries were multiplex-sequenced in duplicate using two flow cells, each containing all 24 libraries to increase the read coverage. The 150-bp paired-end sequence reads were generated by the Illumina NextSeq 500 platform. A total of 89.2 and 94.7 gigabytes of sequencing data were generated from each flow cell, after which the duplicate raw read files for each library generated from these two flow cells were concatenated using Windows Command Prompt. The raw sequence data has been deposited in the NCBI SRA as BioProject PRJNA748697.

### Read mapping and differential gene expression analysis

All reads were trimmed for adapters in CLC Genomics Workbench version 12.0. Quality control of the raw sequence reads was further performed by discarding reads with a Phred score less than 20 and length shorter than 100 bp. After trimming, the average Phred score for all paired-end reads of 24 libraries was 30, ranging from 21 to 36. The trimmed, paired-end reads were mapped using the official gene set version 1.0 (OGSv1.0) [64, 65] of the *Frankliniella occidentalis* genome [35]. The per- and cross-sample library size normalization were performed on the mapped reads by RNA-Seq Analysis tools in CLC Genomic Workbench. Differential gene expression analysis was then completed using the normalized read count data in value of transcripts per million (TPM) for each treatment (V/NV) of corresponding sampling time point-stage (3-L1, 24-L2, and 48-L2) from all four biological replicates. Number of differentially-expressed transcripts and their fold change values were obtained after a cutoff of 2.0 for absolute fold change and 0.05 for false discovery rate (FDR) corrected *p*-values.

### Gene ontology analysis of differentially-expressed transcripts (DETs)

Nucleotide sequences of all DETs identified across the three sampling time points (147 non- redundant) were subjected to Blastx using the NCBI non-redundant protein database and Gene Ontology (GO) functional annotation (E<1.0E-6) using the CloudBlast resource in Blast2GO v5.2.5 [66]. The DETs with GO terms assigned were functionally categorized based on their molecular function, biological process, and subcellular localization [67].

### Validation of differential gene expressions using Droplet Digital PCR

Differential expression of ten selected transcripts in response to virus at 3-L1 stage was examined using Droplet Digital PCR (ddPCR) (Bio-Rad, Hercules, CA). Because of the difficulty in obtaining a large amount of gut RNA samples and the limited sensitivity of conventional qPCR technique to detect gene expression at a low level, the ddPCR was employed for the gene expression assay. This analysis was performed on gut samples obtained from three biological replications of another, independent virus acquisition assay, instead of using the remaining RNA samples from the RNA-Seq experiment. Primers for target DETs were designed using Primer3Plus (http://www.bioinformatics.nl/cgi-bin/primer3plus/primer3plus.cgi). Primer sequences are listed in **Table S5**. The cDNA sample was synthesized from 500 ng of 100-pooled gut RNA and diluted in 16-fold. 8 ul diluted cDNA were added into 12 ul EvaGreen reaction mixture containing 200 mM of sequence-specific primer pairs following the manufacturer’s protocol. The internal reference gene, Actin and water control were included in ddPCR and used for normalizing the estimated copy number of target transcripts. Two technical replicates of 20 ul of EvaGreen reactions for target transcripts, Actin, and water control were included in each ddPCR run. The abundance of the target transcript (copies/ul) was estimated by QuantaSoft software v1.7 (Bio-Rad, Hercules, CA). The log2 fold change of each target transcript in virus-exposed guts against non-exposed guts was calculated after normalization to the reference gene (target gut transcript/reference) [68].

### Weighted gene co-expression network analysis

The pairwise relationships among all thrips transcripts (a total of 16849 transcripts) across RNA-Seq samples were analyzed using the weighted gene co-expression network analysis (WGCNA) [36]. The TPM-normalized read count data for 24 RNA-Seq samples was used for network construction to detect modules (clusters) of highly correlated transcripts, followed by eigengene calculation for each module and relating modules to external sample traits (time-larval stages) using eigengene network. The weighted network analysis was carried out using the WGCNA R software package, following the tutorials with “Step-by-step network construction and module detection” methodology (last updated on February 13, 2016) developed by Langfelder and Horvath (https://horvath.genetics.ucla.edu/html/CoexpressionNetwork/Rpackages/WGCNA/Tutorials/) [36]. Briefly, the input count data was cleaned for excessive missing values, after which all 24 samples were preserved for downstream analyses. The expression samples were matched to their corresponding external traits for time-larval stages (3-L1, 24-L2, and 48-L2). A soft-thresholding power of 12 was selected, of which value reached 0.97 of the scale-free topology fit index. The adjacencies between genes were calculated using the “Signed” network after raising the co-expression similarity values by the power of 12. The hierarchical clustering tree of genes was generated with the topological overlap matrix (TOM)-based dissimilarity. Modules of highly co-expressed genes were identified using the Dynamic Tree Cut method with minimum module size set as 30 and their corresponding eigengenes were calculated. The highly correlated modules were merged into one if the pairwise correlation of their eigengenes was larger than 0.75. The modules that were significantly associated with the external traits (time-larval stages) were identified and quantified by analyzing their corresponding correlations and *p*-values.

Considering the significance of identified module-trait correlations and the number of DETs assigned to each module, the turquoise, blue, and light cyan modules were selected for further analysis (**Table S3**). The DETs identified across the three time-larval stages were exported from each selected module and used for constructing the intramodular interaction networks. The transcripts that fell into the top 10% of both module membership (kME, correlation of a node to a module eigengene) and intramodular connectivity (kIN, equals to the sum of connection weights between a node and all its network neighbors) were considered as the candidate hub transcripts. The intramodular interaction networks of DETs within turquoise, blue, and light cyan modules were visualized using VisANT software platform v5.53 [69]. For better visualization of the interactions, a weight cutoff of ≥0.15 was applied for turquoise and blue networks, and a weight cutoff of ≥0.35 was applied for light cyan network.

### Gene expression analysis of hub and select connecting DETs in larval guts during and after virus exposure

A time-course experiment was performed to analyze the temporal expression of selected hub transcripts and their connecting transcripts (see **Figure S1.2** for guided illustration). The virus acquisition experiment was set up similarly as the RNA-Seq experiment described above, where 24h AAP was given to the synchronized L1s followed by a 24h feeding period on healthy green bean pods. However, a subsample of larvae (∼30-40) from each treatment were sequentially collected after 6h, 18h, 24h, 30h, 42h, and 48h since the first exposure to virus-infected leaves. Larvae from the first three time points were collected during virus exposure (DVE, 24h AAP) and the rest were collected post virus exposure (PVE, feeding on healthy green beans). Twenty dissected gut tissues were pooled per sample and extracted for RNA using PicoPure RNA isolation kit (Arcturus) following the manufacturer’s protocol. 130 ng of RNA were reverse transcribed into cDNA using a Verso cDNA kit and diluted in 4-fold with nuclease-free water. The expression levels of target gut transcripts were detected in ddPCR following the procedure described above. The estimated abundance of target transcript was normalized to the internal reference *F. occidentalis* Actin gene. Additionally, the abundance of TSWV-N and NSs RNA were also estimated at each sampling time points using qPCR as described above and normalized to the internal reference *F. occidentalis* Actin gene using the inverse equation in Pfaffl [70] as described in Rotenberg 2009 [71]. Primer sequences are listed in **Table S5**. The reliability of Actin as an invariant reference gene for normalized expression analysis in gut tissues between treatments within each sampling time point was confirmed with no more than 2.8% of the coefficient of variation in Actin Ct values between virus-exposed and non-exposed gut samples. The time-course experiment was independently repeated three times.

Statistical analysis of temporal expression data was performed by fitting to a generalized linear mixed model using the PROC GLIMMIX procedure of SAS 9.4 (SAS Institute, Cary, NC) with virus condition, time, and their interaction as the three fixed effects. The Gaussian distribution (link function = identity) and restricted maximum likelihood estimation method were specified. The statistically significant effects on gene expression levels were determined by the Type III tests of fixed effects and least square means using the LSMEANS ’Slice’ and ’Slicediff’ options with a Tukey adjustment option for multiple comparisons (*P*<0.05). One-way ANOVA following Tukey’s multiple comparisons (JMP Pro 15.2.0) was used to test the statistical significance in virus abundance between treatments. Pearson’s correlation was performed between log2 transformed TSWV-N or -NSs normalized abundance and log2 transformed expression of target transcripts in JMP Pro 15.2.0.

## DECLARATIONS

### AVAILABILITY OF DATA AND MATERIALS

Raw sequencing data generated and used for this study are available in the NCBI Sequence Read Archive, under Bioproject ID: PRJNA748697.

## ACKNOWLEDGEMENTS

We thank David A. Baltzegar, Director of the Genomic Sciences Laboratory at North Carolina State University, for providing technical advice about RNA sequencing, and we thank Anna E. Whitfield for internal review of the final version of the manuscript.

## AUTHOR INFORMATION

### Affiliations

Department of Entomology and Plant Pathology, North Carolina State University, Raleigh, NC, 27695, United States Jinlong Han & Dorith Rotenberg (corresponding author) Authors’ contributions DR conceptualized the project, JH and DR designed the experiments, JH performed the experiments and analyzed the data; JH drafted the manuscript, and JH and DR revised the manuscript. All authors have read and approved the final manuscript.

## ETHICS DECLARATIONS

Ethics approval and consent to participate Not applicable.

## Consent for publication

Not applicable

## ADDITIONAL FILES

Figure S1. Experimental design illustrations (.docx)

Table S1. Sequence read quality and mapping metrics for each *Frankliniella occidentalis* larval gut RNAseq library (.docx)

Table S2. Differentially-expressed transcripts (DETs) identified in gut tissues of *Frankliniella occidentalis* larvae (3-L1, 24-L2, and 48-L2) in response to TSWV infection (.xlsx)

Table S3. Module memberships of the 147 non-redundant, differentially-expressed transcripts (DETs) distributed across 14 modules generated from a weighted gene co-expression network analysis (WGCNA) (.xlsx)

Table S4. Strength of intramodular membership, level of connectivity, and annotations for the differentially-expressed transcripts (DETs) in the turquoise, blue, and lightcyan module networks (.xlsx)

Table S5. Primer pairs designed for normalized quantification of tomato spotted wilt virus (TSWV) abundance and differentially-expressed transcripts in response to TSWV in *Frankliniella occidentalis* larval guts (.docx)

