## Supplementary material for "Integration of transcriptomics and network analysis reveals co-expressed genes in *Frankliniella occidentalis* guts that respond to tomato spotted wilt virus infection": Figure S1

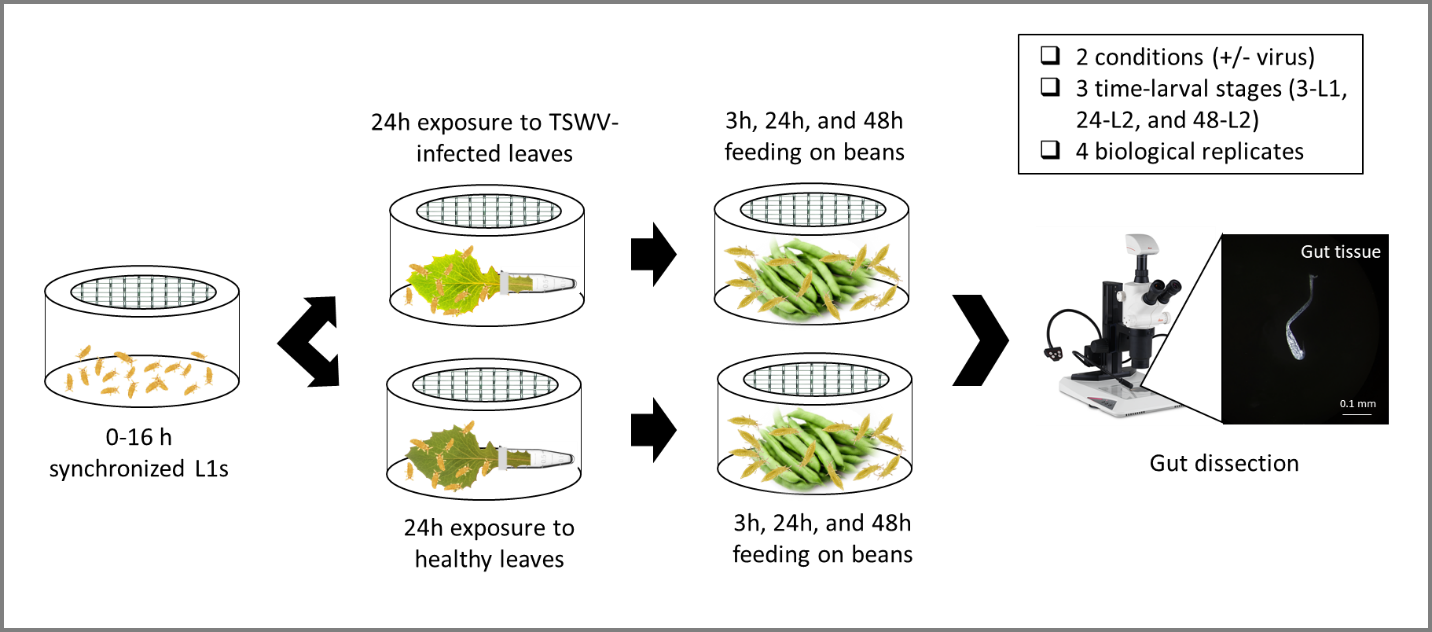

**Figure S1.1.** Experimental design for preparing gut samples for RNA-Seq analysis. Cohorts of first instars (L1) were given a 24-hour acquisition access period on tomato spotted wilt virus-infected plant tissue (or healthy tissue for control group), then virus inoculum was replaced with healthy green bean pods to allow larvae to develop, followed by sub-sampling of larval groups at 3 hours (3-L1), 24 hours (24-L2, early second instar), and 48 hours (48-L2, late second instars) after removal of the virus inoculum. Host plant: *Emilia sonchifolia,* the lilac tasselflower. Number of gut pooled per sample = 100.

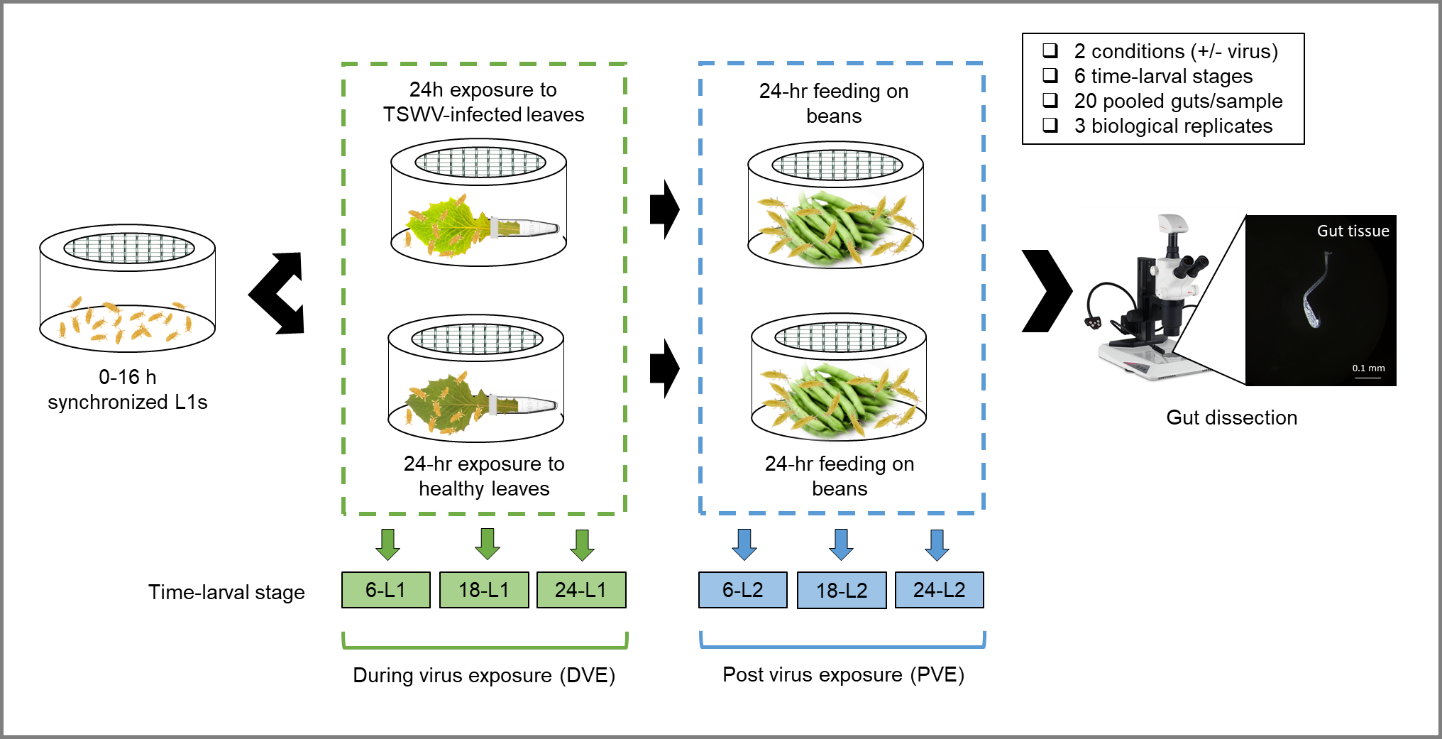

**Figure S1.2.** Illustration of the time-course experiment designed to evaluate gene expression in *Frankliniella occidentalis* larval gut tissues during and after virus exposure. The same virus acquisition assay was performed as described in Figure S1.1, except groups of larvae were collected at three time points during virus exposure (6h, 18h, 24h) and three time points after removal of the virus inoculum (6h, 18h, 24h = 30h, 42h, and 48h after L1s first exposed to virus) virus exposure. L1 = first instar larvae, L2 = second instar larvae.
