## Supplementary material for "Integration of transcriptomics and network analysis reveals co-expressed genes in *Frankliniella occidentalis* guts that respond to tomato spotted wilt virus infection": Table S1

**Table S1**. Sequence read quality and mapping metrics for each *Frankliniella occidentalis* larval gut RNAseq library.

| library | Bio-rep | Raw reads | ^a^Filtered reads | Filtered reads rate (%) | Average length of filtered reads (bp) | ^b^Filtered Q30 reads rate (%) | ^c^Mapped to genes rate (%) |
| --- | --- | --- | --- | --- | --- | --- | --- |
| NV-3-L1 | 1 | 81,642,536 | 65,845,687 | 81 | 143 | 88 | 89 |
|  | 2 | 75,765,982 | 61,289,827 | 81 | 143 | 89 | 87 |
|  | 3 | 84,833,834 | 67,809,089 | 80 | 143 | 89 | 85 |
|  | 4 | 78,501,996 | 62,912,386 | 80 | 143 | 88 | 86 |
|  | Mean | 80,186,087 | 64,464,247 | 80 | 143 | 89 | 87 |
| V-3-L1 | 1 | 84,017,176 | 67,273,979 | 80 | 143 | 89 | 88 |
|  | 2 | 73,729,020 | 59,583,731 | 81 | 143 | 89 | 87 |
|  | 3 | 84,326,476 | 67,793,623 | 80 | 143 | 89 | 86 |
|  | 4 | 72,364,248 | 58,247,239 | 80 | 143 | 88 | 87 |
|  | Mean | 78,609,230 | 63,224,643 | 80 | 143 | 89 | 87 |
| NV-24-L2 | 1 | 63,638,408 | 48,729,955 | 77 | 142 | 88 | 86 |
|  | 2 | 77,267,704 | 62,316,265 | 81 | 143 | 89 | 85 |
|  | 3 | 80,830,084 | 64,561,559 | 80 | 143 | 89 | 87 |
|  | 4 | 76,968,534 | 60,830,273 | 79 | 143 | 89 | 85 |
|  | Mean | 74,676,183 | 59,109,513 | 79 | 143 | 89 | 86 |
| V-24-L2 | 1 | 80,874,870 | 65,283,468 | 81 | 143 | 88 | 86 |
|  | 2 | 75,842,626 | 60,932,174 | 80 | 143 | 89 | 85 |
|  | 3 | 79,632,852 | 63,914,730 | 80 | 143 | 88 | 87 |
|  | 4 | 77,480,454 | 61,874,563 | 80 | 143 | 89 | 86 |
|  | Mean | 78,457,701 | 63,001,234 | 80 | 143 | 89 | 86 |
| NV-48-L2 | 1 | 80,069,000 | 64,333,508 | 80 | 143 | 88 | 85 |
|  | 2 | 77,658,008 | 56,572,753 | 73 | 142 | 87 | 85 |
|  | 3 | 99,665,504 | 79,984,720 | 80 | 143 | 88 | 85 |
|  | 4 | 75,121,230 | 60,864,727 | 81 | 143 | 88 | 87 |
|  | Mean | 83,128,436 | 65,438,927 | 79 | 143 | 88 | 85 |
| V-48-L2 | 1 | 76,486,916 | 60,857,415 | 80 | 143 | 89 | 85 |
|  | 2 | 84,786,430 | 68,428,567 | 81 | 143 | 88 | 86 |
|  | 3 | 79,401,852 | 62,858,294 | 79 | 142 | 88 | 85 |
|  | 4 | 71,096,576 | 56,186,572 | 79 | 143 | 89 | 85 |
|  | Mean | 77,942,944 | 62,082,712 | 80 | 143 | 89 | 85 |

^a^High quality sequence data was obtained after filtering the raw data using trimming tools in CLC Genomics Workbench; ^b^The filtered Q30 reads rate = arithmetic mean of sequence base qualities higher than the Phred score of 30; ^c^The filtered paired end reads were mapped to the official gene set v1.0 (OGSv1.0) of the *Frankliniella occidentalis* genome reference (1, 2).

1. Rotenberg D, Gibbs RA, Worley KC, Murali SC, Lee SL, Muzny DM, Hughes DST, Chao H, Dinh H, Doddapaneni H, Qu J, Dugan-Perez S, Han Y, Richards S: ***Frankliniella occidentalis* genome assembly v1.0. Ag Data Commons (Database).** [https://doi.org/10.15482/USDA.ADC/1503960 2019](https://doi.org/10.15482/USDA.ADC/1503960%202019).

2. Rotenberg D, Robertson HM, Oliver JE, Benoit JB, Snoeck S, Baumann AA, Ben-Mahmoud S, Veenstra JA, Jacobs CGC, Park Y, Panfilio KA, Widana Gamage, S. M. K., Ahn SJ, Vargas Jentzsch IM, Jennings EC, Szuter EM, Christiaens O, Scanlan J, Taning CNT, Martynov A, Bejerman NE, Nair A, Jones JW, Friedrich M, van der Zee M, Dearden P, Minakuchi C, Armisén D, Schneweis DJ, Melo FL, Poelchau M, Dermauw W, Hughes DST, Richards S, Didion EM, Holmes CJ, Rosendale AJ, Rosselot A, Dolan A, Whitfield AE, Ullman DE, Dietzgen RG, Smagghe G, Van Leeuwen T, Warren JH: ***Frankliniella occidentalis* Official Gene Set OGSv1.0. Ag Data Commons (Database).** <https://doi.org/10.15482/USDA.ADC/1504029> 2019.
