## Supplementary material for "Integration of transcriptomics and network analysis reveals co-expressed genes in *Frankliniella occidentalis* guts that respond to tomato spotted wilt virus infection": Table S5

**Table S5.** Primer pairs designed for normalized quantification of tomato spotted wilt virus (TSWV) abundance and differentially-expressed transcripts (FOCC) in response to TSWV in *Frankliniella occidentalis* larval guts

| **Sequence name or transcript ID** | **^a^Primer sequence (5’-3’, forward/reverse)** | **Amplicon length (bp)** |
| --- | --- | --- |
| ^b^TSWV-N | GCTTCCCACCCTTTGATTC/  ATAGCCAAGACAACACTGATC | 140 |
| ^b^TSWV-NSs | ACTCTGTTCTGGCACTATCTG/  GCTGGAATCGGTCTGTAATAT | 147 |
| ^bcd^Actin | GGTATCGTCCTGGACTCTGGTG/  GGGAAGGGCGTAACCTTCA | 69 |
| ^c^FOCC001422 | GGACTCGCCAGCGGATCTGA/  GCTCCCGAAGGTACGCCTGA | 186 |
| ^c^FOCC000537 | TGAACCGCGTTGAAGAAAGC/  AGTAGCCTTAGTTTCCGTCAGG | 190 |
| ^c^FOCC014170 | AGCATGGGTTCTACGTCCGCA/  ACAGTGCTCCCGAAGACCTGG | 138 |
| ^c^FOCC014171 | CCTGGCGGAGACTGAGGCAA/  GGAGCCGCTGTAAGGAAAGGG | 123 |
| ^c^FOCC006994 | CAACCAGACATCAAACGACCC/  CATTACAGGCACACAGAAGTCC | 122 |
| ^c^FOCC012626 | ACCGGAGACAAGCCCACCAT/  GGTTCGAAACCACCGTCGGA | 141 |
| ^c^FOCC003912 | TCCCATGGTGTACCTGCCGT/  TGGCAAGCTACGCGATCACT | 108 |
| ^c^FOCC011985 | AGGTGCGGATTGTTGTTAGTGCT/  TCCTACAAGAAAAAGGAAGTGGCA | 207 |
| ^c^FOCC017299 | CATCTACCACTTCGACCACCC/  GTGCTTGTTGTGTCGTTCATACC | 143 |
| ^c^FOCC014240 | TCCGGGGGTACTCCTGCCTA/  GGCGCATGTCCGGTTTTGGA | 175 |
| ^d^FOCC013675 | CTGTACAGCACCAGCAATGT/  TCCAGGTGTTGTTGGAGTCG | 193 |
| ^d^FOCC004976 | CGGCTGTGAGCAAACCAAAT/  CTTTGAATCCGTCGCCACTT | 163 |
| ^d^FOCC013756 | ACTACAAGGGCGGAATGTGG/  CCGTTGCCAGTCCCGTATTA | 168 |
| ^d^FOCC001422 | AGGACGGCAACAAGGTCATT/  AAGGTCCAGGAGTTCTTCGG | 145 |

^a^Primers were designed using Primer3Plus and Primer-Blast in NCBI.

^b^Primer pairs used in qPCR for testing virus titers in larval guts; ^c^Primer pairs used in ddPCR for validating the RNAseq analysis of virus-responsive larval gut transcripts; ^d^Primer pairs used in ddPCR time-course experiment for evaluating expression levels of hub and connecting transcripts in the blue and turquoise networks.
